# Sensitivity to sequencing depth in single-cell cancer genomics

**DOI:** 10.1101/213744

**Authors:** João M. Alves, David Posada

**Affiliations:** Department of Biochemistry, Genetics and Immunology, University of Vigo, Spain.; Biomedical Research Center (CINBIO), University of Vigo, Spain.; Galicia Sur Health Research Institute, Vigo, Spain.

**Keywords:** single-cell sequencing, intratumor genetic heterogeneity, variant calling, clonal inference, tumor phylogenies.

## Abstract

**Background:** Querying cancer genomes at single-cell resolution is expected to provide a powerful framework to understand in detail the dynamics of cancer evolution. However, given the high costs currently associated with single-cell sequencing, together with the inevitable technical noise arising from single-cell genome amplification, cost-effective strategies that maximize the quality of single-cell data are critically needed. Taking advantage of five published single-cell whole-genome and whole-exome cancer datasets, we studied the impact of sequencing depth and sampling effort towards single-cell variant detection, including structural and driver mutations, genotyping accuracy, clonal inference and phylogenetic reconstruction, using recent tools specifically designed for single-cell data.

**Results:** Altogether, our results suggest that, for relatively large sample sizes (25 or more cells), sequencing single tumor cells at depths >5x does not drastically improve somatic variant discovery, the characterization of clonal genotypes or the estimation of phylogenies from single tumor cells.

**Conclusions:** We demonstrate that sequencing many individual tumor cells at a modest depth represents an effective alternative to explore the mutational landscape and clonal evolutionary patterns of cancer genomes, without the excessively high costs associated with high-coverage genome sequencing.

## Background

Recent advances in next-generation sequencing (NGS) technologies revealed that the large majority of cancer genomes are heterogeneous despite their monoclonal origin, with the continuous expansion of the tumor mass contributing to the accumulation of somatic mutations within malignant cells, hence promoting the proliferation of distinct genetic lineages (i.e., clones) through time [1]. While quantifying this intratumor heterogeneity (ITH) remains a difficult task, as standard methods in cancer genomics generally rely on population-level analysis from bulk experiments, single-cell sequencing (hereon SC-Seq) approaches are now widely viewed as a promising alternative to explore tumor evolution [2]. Indeed, a collection of recent studies have successfully applied SC-Seq to determine the mutational load in individual tumors [3], estimate the frequency of subclones [4], infer evolutionary relationships [5], or explore the role of ITH in metastatic dissemination [6].

Nevertheless, several technical challenges surrounding current SC-Seq methodologies greatly limit our ability to obtain reliable genomic information from single cells. For instance, the multiple rounds of whole genome amplification (WGA) usually required prior to SC-Seq are known to introduce a high number of sequence artifacts that can be confounded with genuine biological variation (see [7] for a detailed review). Other technical errors, such as insufficient physical coverage, uneven genome amplification, and allelic dropout, may also generate substantial artificial variability in cancer genomes, compromising the ability to detect real somatic heterogeneity from SC-Seq data [8]. As a consequence, alternative strategies are needed in order to eliminate the noise generated during WGA while effectively allowing the quantification of ITH from single cells.

Zhang et al. (2015) [9] started addressing some of these issues and demonstrated the efficiency of a census-based strategy for accurate variant detection in single-cells. By using multiple cells and trusting only variants detected in at least two single-cell libraries, they detected up to 80% of germline SNPs in the human chromosome 5 with 59 cells sequenced at 0.3x or 22 cells at 1x. Their results suggest that for detecting clonal and subclonal variants in single-cells and given a fixed sequencing effort, it is best to sequence multiple cells (in their case a minimum of 20) at a modest depth (~1x).

Here, we further explore the sensitivity of SC-seq to sequencing depth using five publicly available single-cell whole-genome (scWGS) and whole-exome (scWXS) cancer datasets. Not only do we expand on the scale of the datasets, but also on the scope of the inferences, including copy-number variant detection, clonal inference and phylogenetic estimation. Altogether, our results suggest that, even though sequencing depth does indeed contribute to a better refinement of somatic variant characterization from tumor single cells, sample size plays a more determinant role for a reliable assessment of the general patterns of somatic variation in cancer genomes. For relatively large sample sizes (e.g., ≧25 samples), sequencing single cells at modest depths (i.e., 5x) enables a similar description of somatic variation, clonal composition, and evolutionary history compared to sequencing depths one order of magnitude higher.

## Results

### Genome coverage

Genome coverage (% of the reference genome covered by ≥1 read) for the single-cell down-sampled datasets decreased non-linearly with lower sequencing depths, in particular when moving from 5x to 1x (**figure 1**).

**Figure 1.**
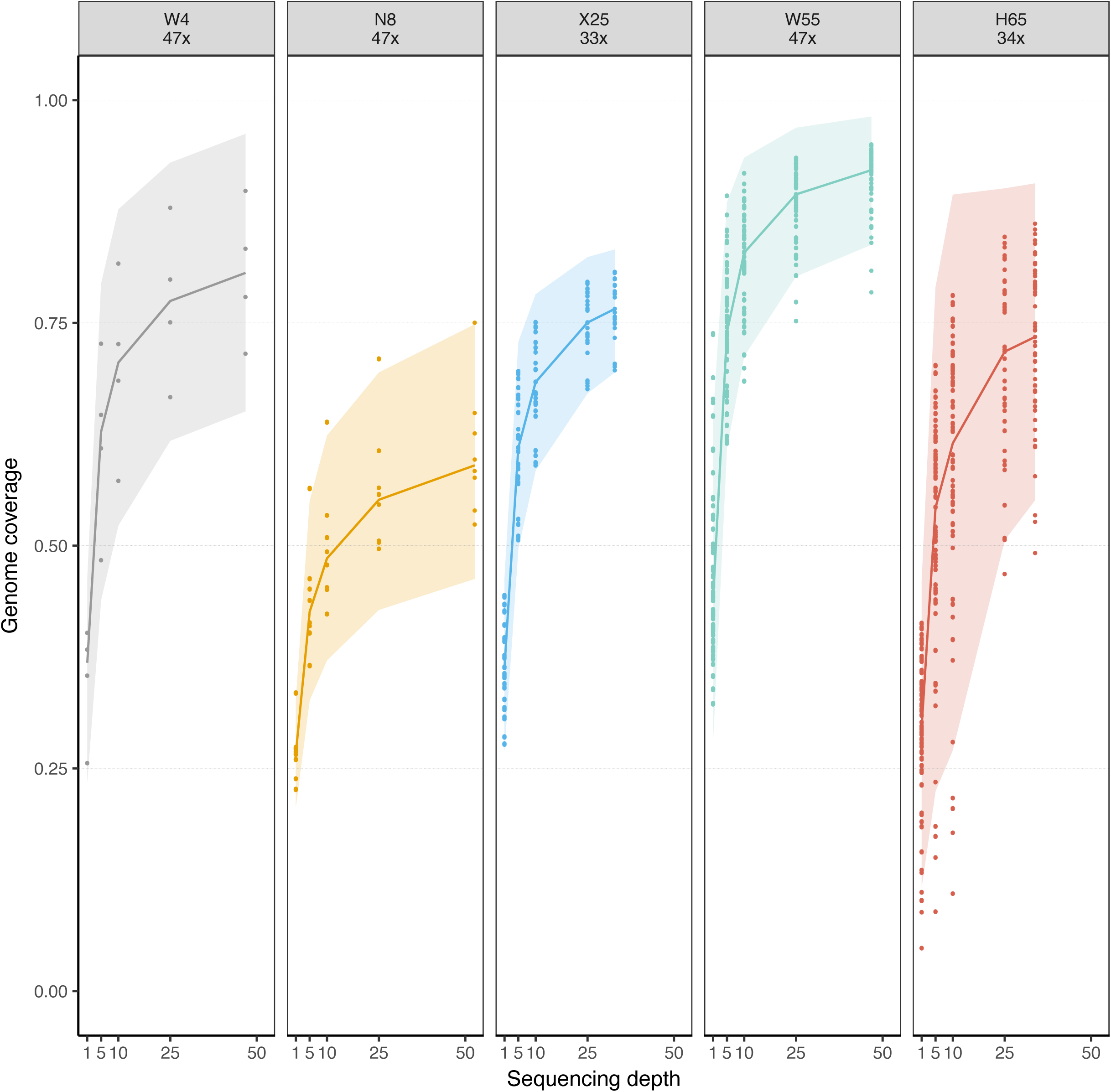
Genome coverage and sequencing depth in the down-sampled single-cell datasets. Each panel depicts a single-cell dataset (e.g., W4) and its original sequencing depth (e.g., 47x). Solid lines represent the average genome coverage obtained for the different replicates at different down-sampled depths. Dots correspond to single cells. Shaded areas illustrate the standard deviation from the mean.

### Single-nucleotide variants (SNVs)

#### SNV detection

The observed decline in genome coverage was logically reflected in the proportion of bulk germline and somatic SNVs found in the single-cell down-sampled datasets (“germline and somatic recall”), which decreased significantly (Tukey HSD p-value < 0.05) at lower sequencing depths (**figure 2**). The germline recall decrease was much less pronounced when the number of cells was large (≥25). Thus, for the X25, W55 and H65 datasets the germline recall was at 1x as high as 70–80% and at 5x was close to 100% (**figure 2A**). On the other hand, when only four or eight cells were available (W4, N8 datasets), the germline recall at 1x decreased dramatically to 5–13%. The somatic recall rate was, as expected, much more modest than for the germline variants (**figure 2B**). The effect of sequencing depth was significant in practically every case. On the other hand, the fraction of SNVs found in the down-sampled replicates that were also identified in the original single-cell datasets (“somatic precision”) (**figure 2C**) was much less affected by sequencing depth, with many non-significant variations between “contiguous” levels of coverage (i.e., 1–5, 5–10, 10–25x).

**Figure 2.**
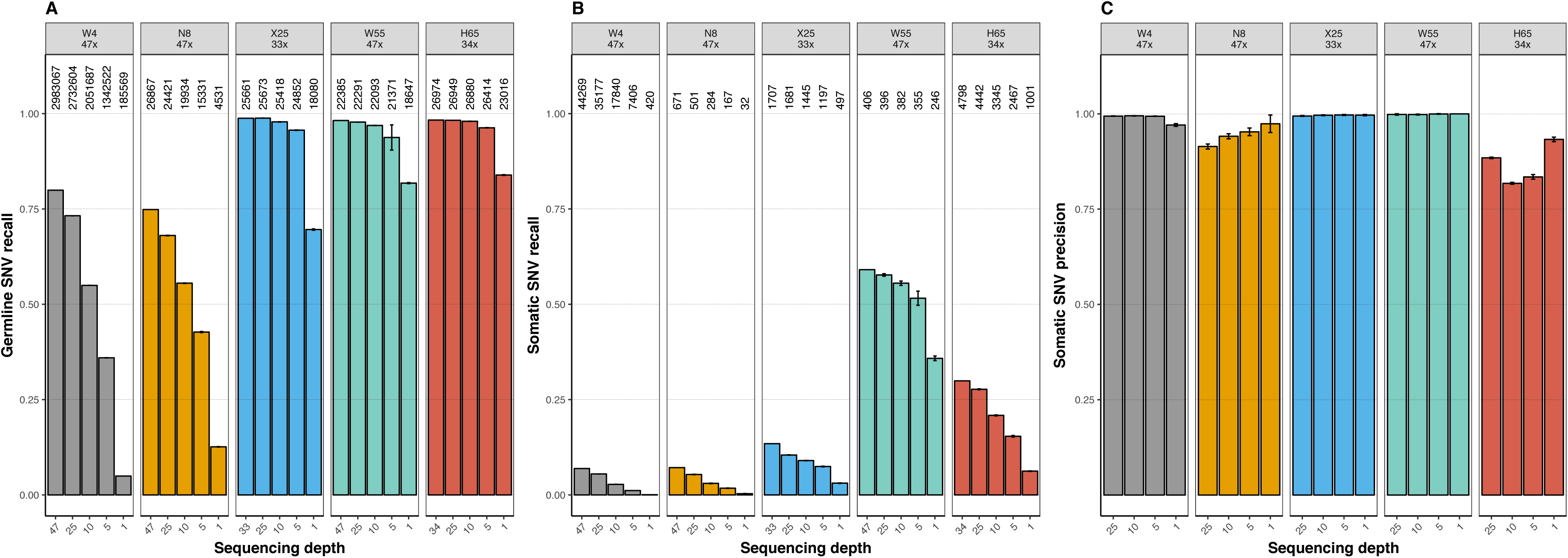
SNV recall and precision in single-cells. (A) Barplots illustrating the proportion of bulk germline variants called in the single-cell down-sampled datasets (germline recall). (B) Proportion of bulk somatic variants identified in the single-cell down-sampled datasets (somatic recall). (C) Proportion of somatic variants called in the down-sampled datasets that were also identified in the original single-cell dataset (somatic precision). Error bars indicate 95% confidence intervals. Numbers above bars indicate number of calls (for B and C these numbers are the same).

#### COSMIC and non-synonymous SNV detection

Interestingly, the somatic recall specific for COSMIC somatic variants (**figure 3 A-B**) decreased very rapidly and significantly (p-value < 0.05) with lower sequencing depths for the smallest datasets (W4 and N8), but not as abruptly for the larger ones, in particular for the X25 and W55 datasets. For example, for the latter the recall was already around 70% at only 5x. A statistically significant trend was also observed for non-synonymous SNVs, which were very difficult to detect at 5x or 1x only for the smaller datasets (**figure 3 C-D**). For larger sample sizes, the non-synonymous SNV recall rate was already above 70% at 1x.

**Figure 3.**
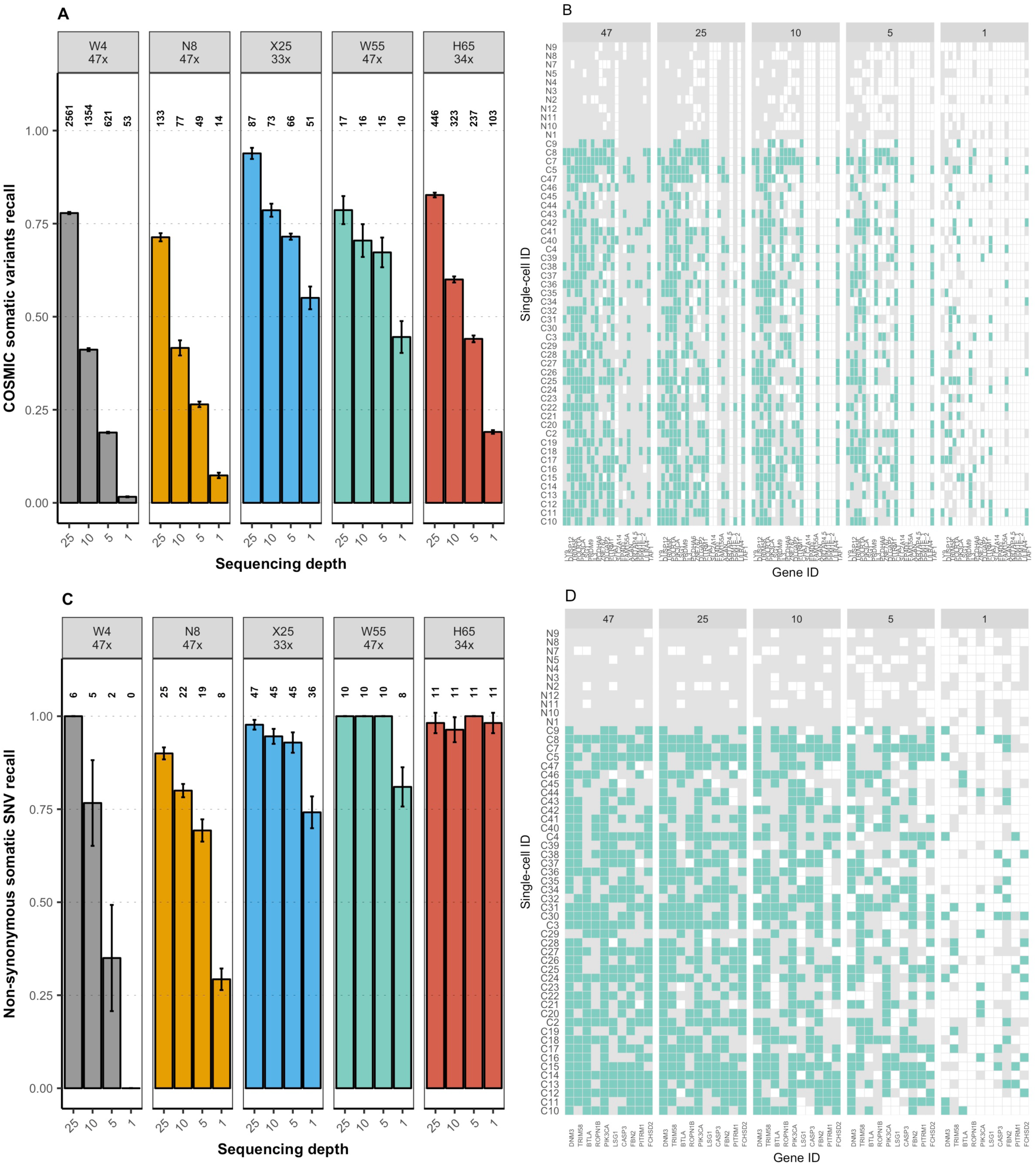
COSMIC and non-synonymous somatic SNV recall in single-cells. (A) Barplots indicate the proportion of bulk COSMIC somatic variants detected in the single-cell datasets (COSMIC recall). Error bars indicate 95% confidence intervals. Numbers above bars indicate number of variants called. (B) Presence-absence profile of COSMIC SNVs across cells for the W55 dataset for replicate 1. Colors illustrate mutation status: Mutated allele - green, Reference allele - grey, Missing data - white. (C) Barplots indicate the proportion of bulk non-synonymous somatic variants detected in the single-cell datasets (non-synonymous recall). (D) Presence-absence profile of non-synonymous SNVs across cells for the W55 dataset for replicate 1. Colors illustrate mutation status: Mutated allele - green, Reference allele - grey, Missing data - white.

#### SNV genotyping

The recall for single-cell SNV genotyping also dropped significantly at lower sequencing depth for all datasets (**figure 4**). Nevertheless, at 5x, 60–90% of the genotypes identified in the original single-cell datasets were already recovered without error. Importantly, discordant SNV genotypes calls were relatively infrequent, and differences between depth levels and datasets were usually due to different amounts of missing calls.

**Figure 4.**
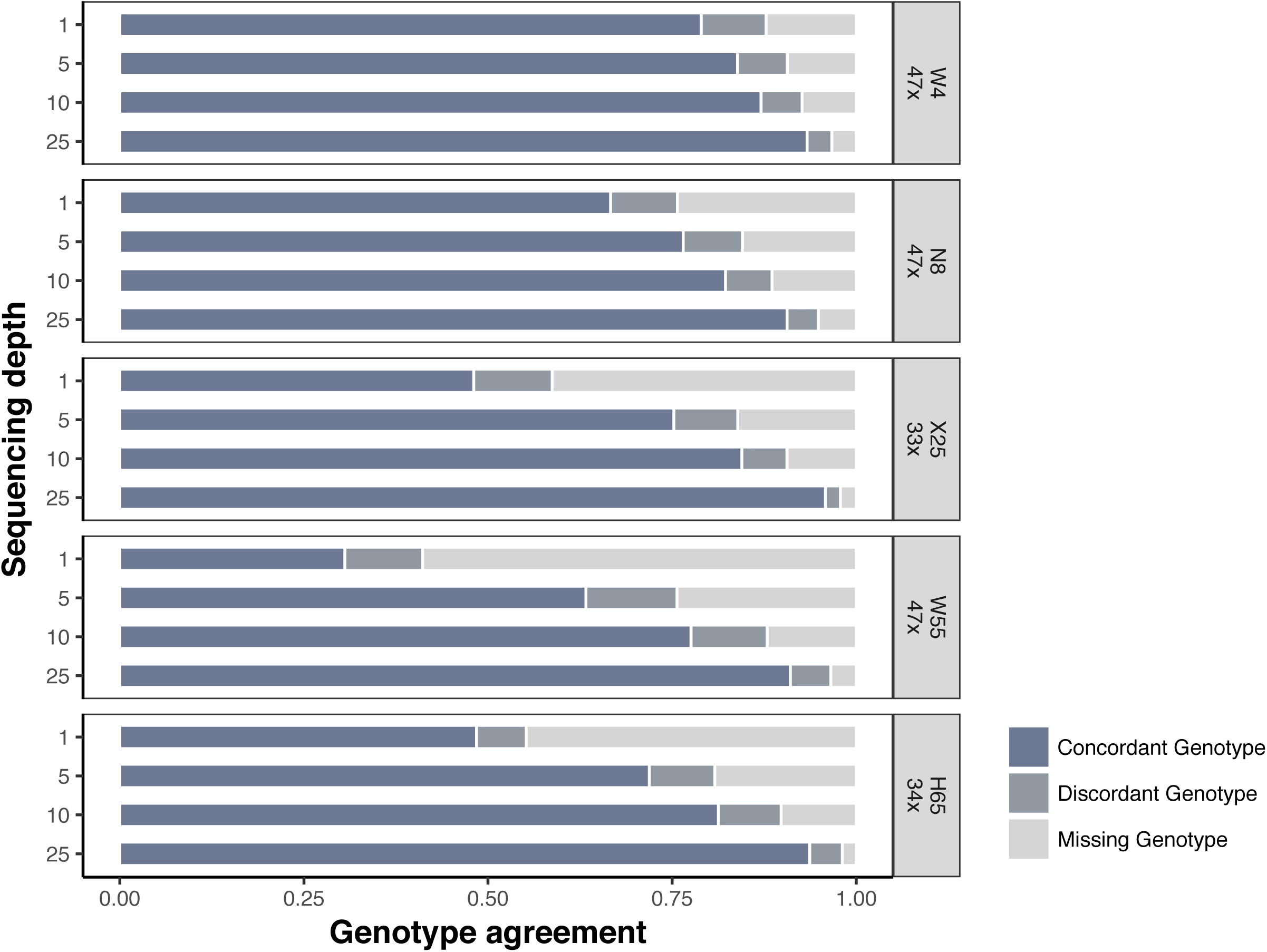
SNV genotype recall in single-cells. Horizontal bars represent the proportion of concordant (dark blue), discordant (dark gray) and missing (light gray) SNV genotype calls (homozygous for the reference allele, heterozygous or homozygous for the alternate allele) for the down-sampled datasets.

### Copy-number variants (CNVs)

Single-cell copy-number profiles were remarkably consistent across sequencing depth (**figure 5**). Breakpoint detection was slightly better for higher sequencing depths, but always quite accurate. For example, more than 70% of the CNV breakpoints inferred from the original dataset were already detected at 1x in all datasets. Moreover, CNV genotype calls were not affected by sequencing depth. Indeed, at 1x the CNV genotype recall was already >99% for all datasets.

**Figure 5.**
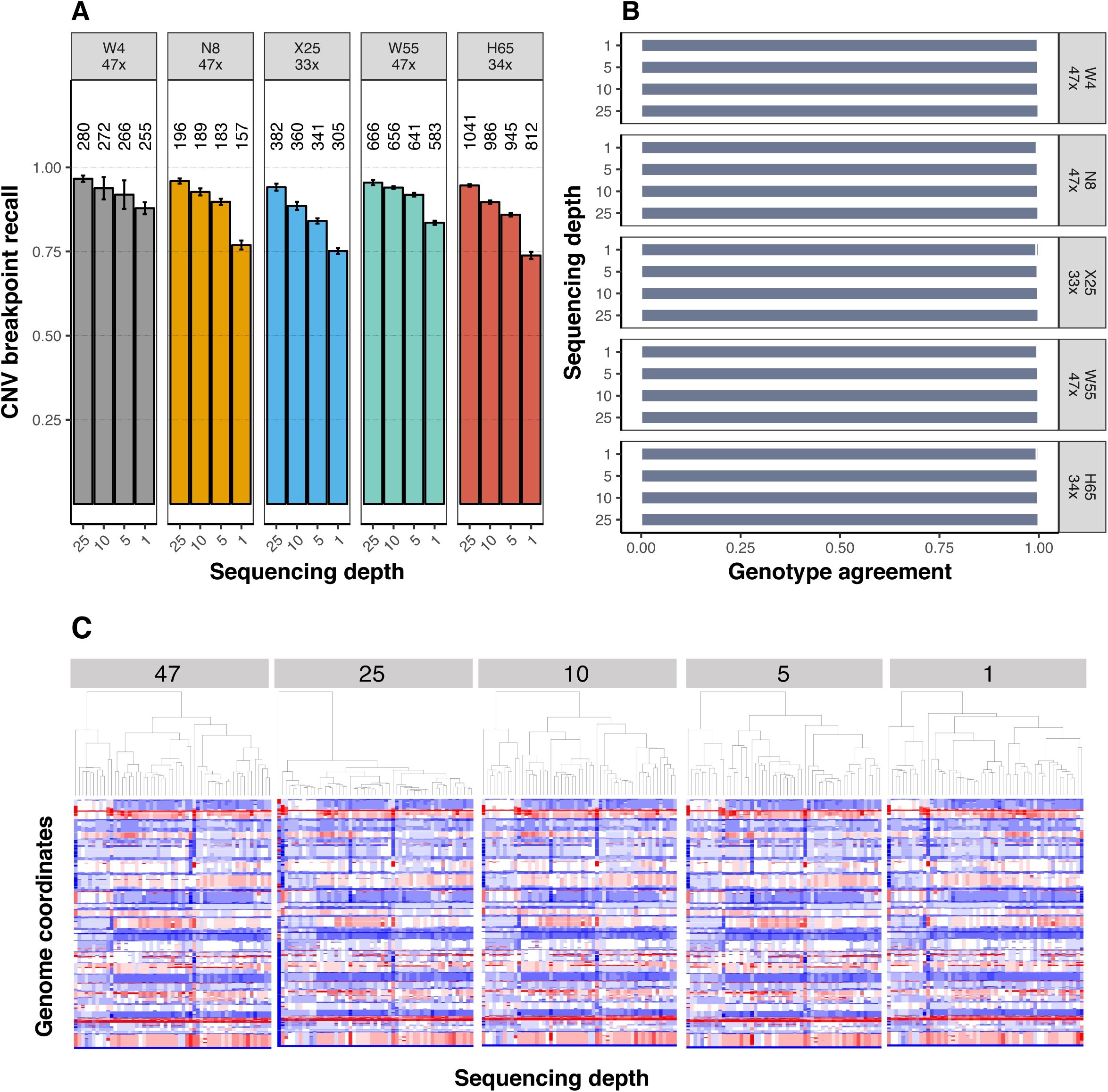
CNV recall in single-cells. (A) Barplots indicate the proportion of CNV breakpoints detected in the down-sampled datasets that were also called in the original single-cell dataset (CNV breakpoint recall). Numbers above bars indicate number of breakpoints detected. Error bars indicate 95% confidence intervals. (B) Horizontal bars represent the proportion of concordant (dark blue), discordant (dark gray) and missing (light gray) CNV genotype calls. (C) Copy-number profiles at different depths for the W55 dataset - replicate 1. Distinct colors represent the CN configuration: CN gain - red, CN loss - blue.

### Clonal genotypes

Clonal inference recall by SGC [21], as measured by the adjusted Rand Index, was not affected by sequencing depth in the smallest and largest datasets (W4, N8 and H65) where the number of inferred clones was always one, but decreased to a different extent at lower depths in the X25 and W55 datasets (**figure 6 A-B**). Indeed, despite the improvements observed at sequencing depths beyond 5x for the X25 dataset, the distinct clonal clusters of the W55 dataset were only distinguishable at 25x.

**Figure 6.**
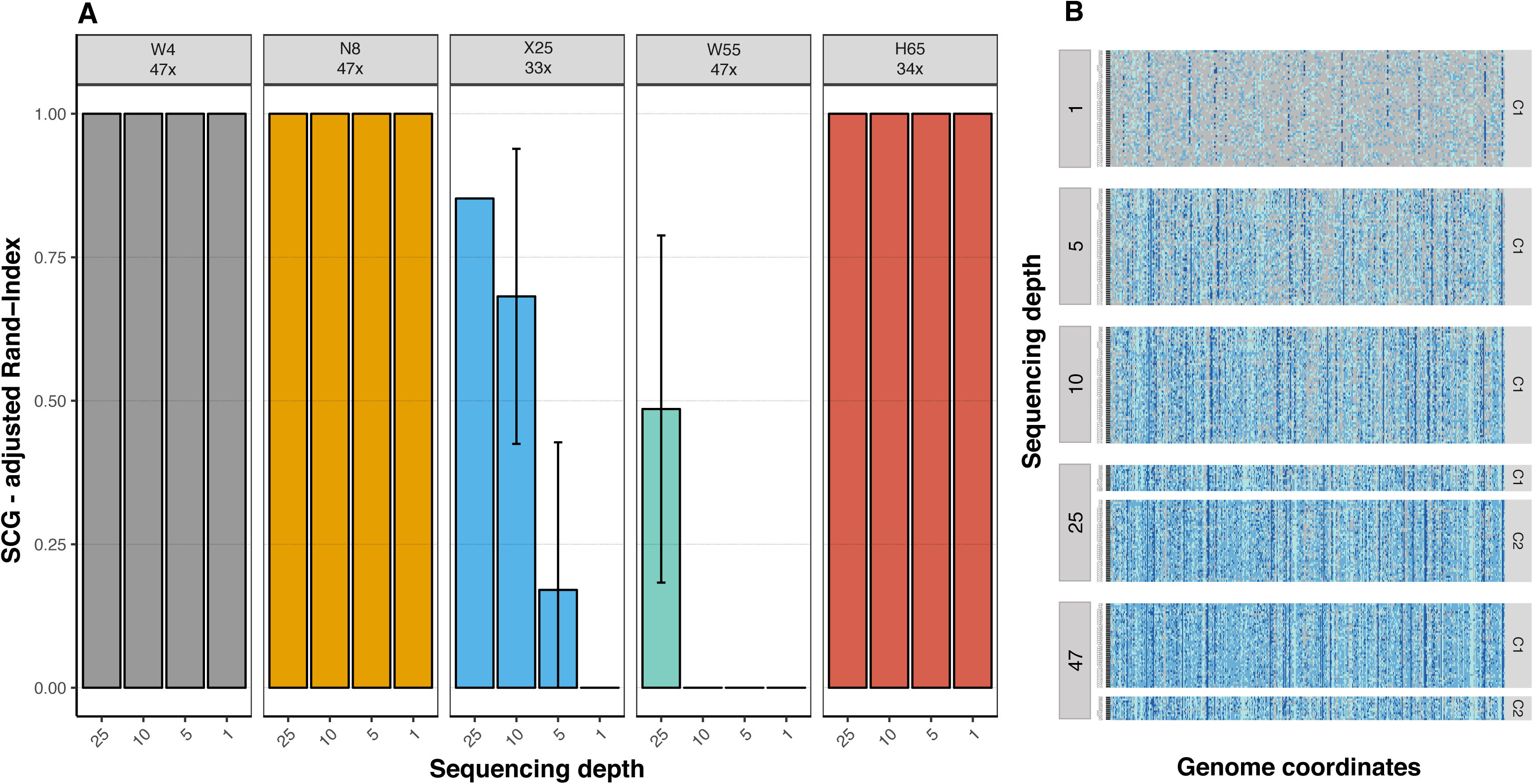
Clonal inference recall from single-cells. (A) Barplots show the adjusted Rand-Index between the clonal clusters inferred from the original dataset and the clonal clusters from the down-sampling experiments. Error bars indicate 95% confidence intervals. (B) SCG clonal clusters at different sequencing depths for the W55 dataset - replicate 1. Genotypes for each locus highlighted in different colors: Homozygous Reference - light blue, Heterozygous - blue, Homozygous Alternative - dark blue, Missing - light gray).

### Clonal trees

In contrast to the smallest datasets, the recall of the clonal trees inferred by OncoNEM [24] (**figure 7 A-B**) was maintained or decreased slightly - not-significantly in multiple occasions - at lower sequencing depths for the larger datasets (X25, N55 and H65) where the number of potential phylogenetic solutions is much bigger.

**Figure 7.**
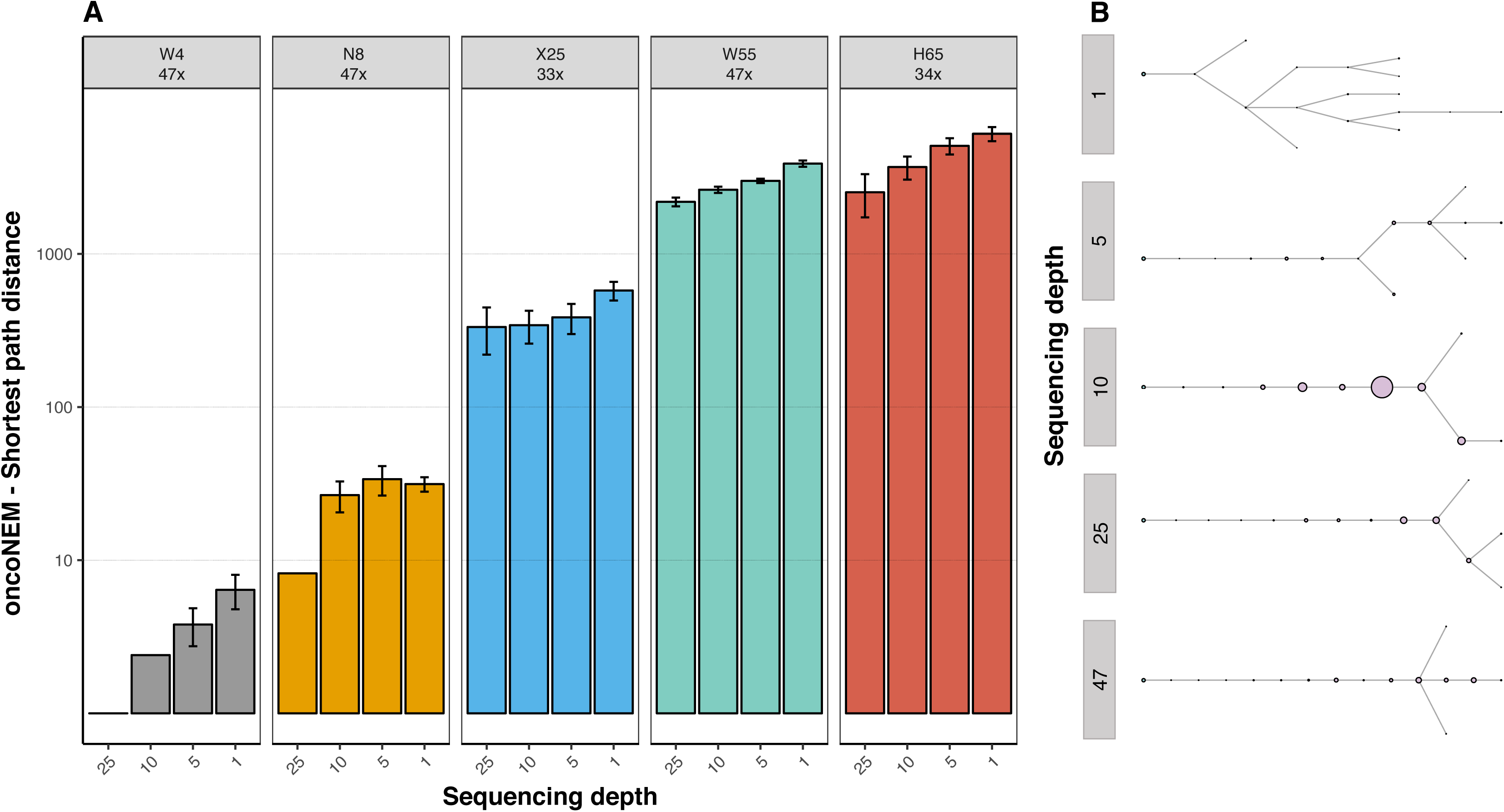
Clonal tree recall from single cells. (A) Barplots show the pairwise cell shortest-path distance between the OncoNEM clonal tree inferred from the original dataset and the OncoNEM clonal trees inferred from the down-sampling experiments. Error bars indicate 95% confidence intervals. (B) OncoNEM clonal trees at different depths for the W55 dataset - replicate 1.

### Single-cell phylogenies

SiFit [25] single-cell phylogenies were also very consistent at sequencing depths equal or larger than 5x (**figure 8 A-B**). In most instances the differences due to depth were not statistically significant. At 1x, in some cases the inferred phylogeny displayed healthy cells intermixed with tumor cells, likely due to poor resolution. Nevertheless, this effect disappeared at 5x, where all tumor cells always clustered together in a single clade, as expected.

**Figure 8.**
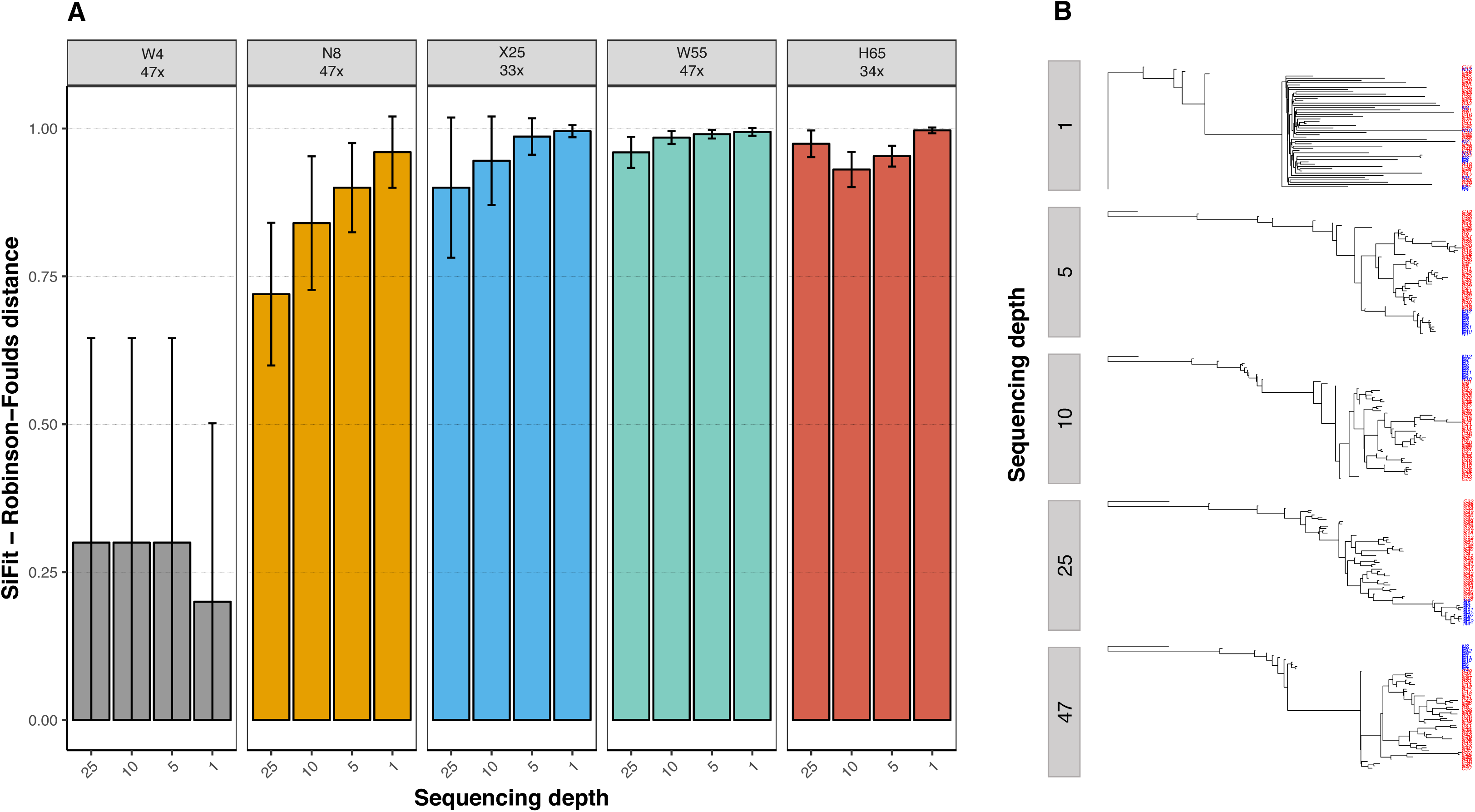
Single-cell tree recall. Barplots show the Robinson-Foulds distance between the SiFit single-cell tree inferred from the original dataset and the SiFit single-cell trees inferred from the down-sampling experiments. Error bars indicate 95% confidence intervals. (B) SiFit unrooted single-cell trees at different depths for the W55 dataset - replicate 1. Distinct colors represent cell type: Healthy cell - blue, Tumor cell - red.

## Discussion

In this study, we aimed to characterize the impact of sequencing depth in single-cell cancer genomics studies. Undeniably, here we have used five datasets with specific characteristics like number of mutations, number of clones, tissue of origin, genomic target, sequencing depth, or amplification bias. In consequence, although some general patterns seem to be more or less clear, care must be taken in generalizing our findings as particular trends may vary for other cancer datasets.

With this caveat in mind, our down-sampling experiments suggest that, overall, larger sequencing depths for small numbers of cells (eight or less) might lead to relevant improvements. In contrast, for relatively large datasets (25 or more cells), our results indicate that sequencing single cells at moderate depths (i.e., 5x) should represent a reasonable approach to characterize the genomic diversity and evolution of tumors, including the identification of putative driver alterations. This is in line with the results of Zhang *et al.* (2015) [9], who showed that for variant detection it is better to have multiple cells sequenced at low depth, given a fixed sequencing effort.

Unsurprisingly, all recalls (SNVs, CNVs, clones, phylogenies) showed some kind of decrease at smaller sequencing depths. In many cases the drop was statistically significant despite being of small magnitude. Notably, for the larger datasets (and by large here we mean – only – dozens of cells), the impact of sequencing depth was much smaller, with the exception of the H65 dataset. This particular dataset, albeit being the largest, displays a very heterogeneous genome coverage for the single cells sampled which may have mislead some of the analyses. Indeed, genome coverage bias has been shown to contribute to a lower sensitivity to detect variants [9], hence potentially explaining some of the somewhat discordant results of the H65 dataset.

In any case, bulk germline SNVs were relatively easy to identify for the three largest datasets even at low sequencing depth. This was indeed expected since germline variants should be present in the vast majority, if not all, of tumor cells. Nevertheless, when the number of single cells was small, the effect of sequencing depth on germline SNV recall was much more pronounced and reached a limit of ~75% at the highest sequencing depth (i.e., 47x) reinforcing the idea that, due to the inherent bias in single-cell genome amplification, broader sampling effort should be favoured over increased sequencing depth in variant detection analysis [9].

While somatic SNVs were much more difficult to detect, it should be highlighted that the number of somatic mutations detected at 5x were usually at the same order of magnitude than the number of mutations detected at higher sequencing depths, except for the smaller datasets. Still, for the smallest dataset analysed (W4), the high number of somatic SNVs detected at 5x (7406) seem plenty enough to conduct many subsequent analyses, like clonal inference or phylogeny reconstruction.

Remarkably, the somatic single-cell SNV precision was, in general, very robust to sequencing depth, suggesting that lower depths do not result in new calls that would not have been made at higher depths. Intuitively, this observation makes perfect sense since at lower sequencing depths the variants detected tend to be the clonal ones (i.e., variants shared by the majority of the single-cells sampled) whereas the detection of low-frequency mutations required higher read depths (data not shown).

One might be worried, however, about missing putative driver mutations, but our results suggest that, as far as the number of single-cells is reasonably large (here 25 or more), most COSMIC somatic variants can be detected at modest sequencing depths (here 5x or more). Similar results were also observed for the somatic non-synonymous variants, suggesting that, in principle, many relevant variants in single-cell genomes are likely to be detected at modest sequencing depths.

Obviously, assigning particular genotypes to the individual cells is a much more involved task than just detecting variants. Importantly, for SNV genotyping, reducing sequencing depth generally resulted in an increased amount of missing data in the single-cell genotype matrix, rather than different genotype calls.

Moreover, and in agreement with previous studies [20,27], CNV characterization from single cells was very robust to sequencing depth, with all down-sampled datasets showing remarkable preservation of CNV breakpoints. Furthermore, CNV genotype assignment was insensitive to the variation in the sequencing depths explored.

It is relatively well established that an accurate identification of clonal genotypes can be very important to understand tumor dynamics and genomic architecture [28–30]. For the datasets analysed here, our results suggest that SC-Seq depth does not affect the identification of tumor clones when the genomic variability between malignant cells is small (i.e., displaying limited clonal population genetic diversification). However, the same was not true for tumors comprising a larger number of subclones, where the different clonal genotypes were only distinguishable at higher sequencing depths. While these results are not necessarily surprising, as clonal identification remains a complex problem even for bulk sequencing data [31,32], they seem to suggest that higher sequencing coverage is ultimately required to resolve fine-scale clonal structure in more heterogeneous tumors.

Finally, in our evolutionary analyses, we observed a moderate impact of sequencing depth with respect to the estimated phylogenetic relationships of the inferred clones and single cells. Perhaps due to the uncertainty stemming from significant amounts of missing data, datasets down-sampled to 1x resulted in phylogenetic trees with healthy cells intermingled with tumor cells, which can be safely considered as artifacts. Otherwise, tree topologies at 5x seemed quite similar to those inferred at higher depths, suggesting that relatively few clonal variants might be enough to resolve the topology of the single-cell trees. Note that the topology does not include branch lengths, whose accurate estimation might require higher sequencing depths.

## Conclusions

In summary, while recent improvements in single-cell isolation, and WGA methods, have led to the development of innovative experimental [9,33–35] and analytical tools [18,20,21,25,35] that reduce the technical noise from SC-Seq data, the costs associated with sequencing multiple single-cell genomes or exomes at high depths are still largely prohibitive. Our results support the idea that sequencing many individual tumor cells at a modest depth, such as 5x, may help circumvent this limitation at least for the type of analyses implemented here. For cancer genomics research, future SC-Seq experiments are likely to benefit from lower sequencing costs, thus setting a higher limit to the amount of SC-Seq data generated. In consequence, more accurate inferences of the clonal architecture of tumor samples are expected to be obtained, which shall ultimately prove crucial to compare models of tumor evolution, trace cell lineages, measure mutation rates, and decipher cell clones responsible for metastatic dissemination and drug resistance [2,36]. Finally, the results obtained here might be extrapolable to some extent to non-tumor single-cell genomes.

## Methods

Five publicly available sequencing datasets from four single-cell studies were retrieved from the Sequence Read Archive (SRA) in FASTQ format, including four single-cell genomes from a breast cancer patient [5] (we will call this dataset “W4” to indicate the authors and the number of cells), eight single-cell exomes from circulating tumor cells from one lung adenocarcinoma patient [10] (“N8” dataset), 25 single-cell exomes derived from a kidney tumor patient [11] (“X25” dataset), 55 single-cell exomes from a breast cancer patient [5] (“W55” dataset), and 65 single-cell exomes from a single JAK-2 negative neoplasm myeloproliferative patient [12] (“H65” dataset). Normal and tumor bulk WGS/WXS data from the same patients were also retrieved. Normal single cells were only available for the three largest datasets. A list of the individual samples and corresponding accession codes is available in **table S1**.

All the analyses enumerated below are described in detail in the accompanying **supplementary note**, including command lines. Both single-cell and bulk reads were aligned to human reference GRCh37 using the *MEM* algorithm in the BWA software [13]. Following a standardized best-practices pipeline [14], mapped reads from all datasets were independently processed by filtering reads displaying low mapping-quality, performing local realignment around indels, and removing PCR duplicates. Raw single-nucleotide variant (SNV) calls for the bulk datasets were obtained using the paired-sample variant-calling approach implemented in the VarDict software [15]. For the N8 dataset, since samples from both primary tumor and metastasis were available, VarDict was run twice, independently for both samples, and the resulting SNVs subsequently merged using the *CombineVariants* tool from the Genome Analysis Toolkit (GATK) [16]. Low-quality SNV calls were removed using the *SelectVariants* tool from GATK. The remaining SNVs were further subdivided into two distinct categories: "germline" SNVs if present in both tumor and normal bulk samples, and "somatic" SNVs if found solely in the tumor bulk samples. Small indels and other complex structural rearrangements were ignored in order to generate a final list of "gold-standard" bulk SNVs. All analyses presented here were based on this set of variants.

The single-cell BAM files were independently downscaled to 25, 10, 5, and 1x sequencing depth using Picard [17]. For each depth level, ten technical replicates were generated for statistical validation, resulting in a total of 6280 BAM files. Single-cell SNV calls were obtained from the original and down-sampled single-cell BAM files using Monovar [18], a variant caller specifically designed for single-cell data, under default settings. Single-cell variant-calling performance was evaluated by estimating the proportion of "gold-standard" germline and somatic bulk SNVs identified in the down-sampled single-cell datasets (germline and somatic recall, respectively). To further characterize the effect of sequencing depth on single-cell variant calling, we determined the fraction of somatic SNVs found in the down-sampled single-cell replicates that were also identified in the original single-cell datasets (“somatic precision”). In addition, we repeated the recall analysis focusing only on the somatic SNVs already described in the Catalogue Of Somatic Mutations In Cancer (COSMIC) database [19] and on the non-synonymous SNVs previously detected (**table S2**).

Single-cell copy-number variants (CNVs) were identified with Ginkgo [20] using variable-length bins of around 500kb. After binning, data for each cell was normalized and segmented using default parameters. Sensitivity was evaluated by assessing the recall of the CNVs and segment breakpoints at the different sequencing depths.

Clonal genotypes were estimated from the somatic SNVs using the Single-Cell Genotyper (SCG) [21] (**supplementary note**), and their recall across sequencing depth was measured with the adjusted Rand-Index (ARI) [22], a version of the Rand-Index corrected for chance [23]. The Rand-Index is a popular statistical measure of the similarity between two data clusterings (corresponding here to groups of mutations, or clones). In addition, clonal trees were also inferred from the somatic SNVs with OncoNEM [24]. Using a similar approach to Ross et al. (2016) [24], the pairwise cell shortest-path distance was used to measure the consistency in tree reconstruction across the different sequencing depths. Furthermore, maximum-likelihood single-cell phylogenies were estimated from the SNVs using SiFit [25]. In this case, phylogenetic recall across sequencing depth was measured using the standard Robinson-Foulds tree distance [26].

Statistical significance for the differences in recall for the experiments described above were assessed using Tukey’s HSD test with a family-wise error rate of 0.05 in R. See the **supplementary note** for a detailed description.

## Declarations

### Ethics approval and consent to participate

Not applicable.

### Consent for publication

Not applicable.

### Availability of data and material

The datasets analysed during the current study are available in NCBI Sequence Read Archive under the following accession numbers: SRA050201(Xu et al., 2012), SRA050202 (Hou et al. 2012), SRP029757 (Ni et al., 2013) and SRA053195 (Wang et al., 2014).

### Author’s contributions

D.P. conceived the project and designed the analyses. J.M.A. performed the analysis. J.M.A. and D.P. wrote the manuscript.

#### Acknowledgments

We would like to thank Sereina Rutschmann, Harald Detering, Laura Tomás, and Sara Rocha for their comments on earlier versions of the manuscript.

